# Accumbal cholinergic interneurons regulate decision making or motor impulsivity depending on latent task state

**DOI:** 10.1101/2025.03.17.643719

**Authors:** Tristan J. Hynes, Chloe S. Chernoff, Kelly Hrelja, Andrew Li, Graeme D. Betts, Lucas S. Calderhead, Catharine A. Winstanley

**Author notes:** Current address: Department of Psychology, University of Cambridge (Tristan J. Hynes, Chloe S. Chernoff). These two authors made equal contributions (Tristan J. Hynes, Chloe S. Chernoff).

## Abstract

Dopaminergic transmission within the nucleus accumbens is broadly implicated in risk/reward decision making and impulse control, and the rat gambling task (rGT) measures both behaviours concurrently. While the resulting indices of risky choice and impulsivity correlate at the population level, dopaminergic manipulations rarely impact both behaviours uniformly, with changes in choice more likely when dopaminergic transmission is altered during task acquisition. Although the task structure of the rGT remains constant, the importance of accumbal dopamine signals relevant for reward prediction versus impulse control may vary over time; the former should dominate while learning which option maximises sugar pellet profits, while the suppression of premature responses becomes more valuable once a decision-making strategy is set and can be exploited. Cholinergic interneurons (CINs) critically control dopamine release within the striatum, and can also encode latent task states deciphered by the frontal cortex. We theorised that aCINs may set the dopaminergic tone of the accumbens to maximise reward learning or impulse control during task acquisition or performance, respectively. Using chemogenetics, we found some support for this hypothesis: activation and inhibition of aCINs once behaviour was stable increased and decreased motor impulsivity in both sexes but had no effect on choice patterns. In contrast, activating and inhibiting aCINs throughout task acquisition did not alter motor impulsivity, but decreased and increased risky choice respectively. However, the former effect was only seen in males and the latter in females. We conclude by proposing a set of testable predictions regarding interactions between acetylcholine and dopamine that could explain these sex differences.

## Introduction

Probabilistic decision making (Phillips *et al*., 2007; Onge *et al*., 2012; Stopper *et al*., 2013) and impulsivity (Dalley *et al*., 2007; Besson *et al*., 2010) are influenced by dopaminergic processes in the nucleus accumbens (NAc). The rat gambling task (rGT) measures both decision making and impulsivity at the same time. Using cell type- and circuit-defined chemogenetic approaches, we have shown that midbrain dopamine neurons (Hynes *et al*., 2021; Hynes *et al*., 2024b) and their projections to the ventral striatum (Hynes *et al*., 2020) causally affect decision making and impulsivity on this task, albeit differentially between males and females, and rarely in concert. Chronic administration of the dopamine D_2/3_ receptor agonist ropinirole also dramatically increased risky choice when administered while animals were still learning the reinforcement contingencies associated with each option, and when reward delivery was paired with audiovisual cues (Mortazavi *et al*., 2023). However, this manipulation had no effect on choice strategies once preferences were well established (Tremblay *et al*., 2017). Conversely, increases in premature responding were more long-lasting and pronounced when the drug was administered to animals that had already attained a stable, asymptotic behavioural baseline, rather than during task acquisition. These data suggest that the degree to which mesolimbic dopamine signals causally regulate decision-making versus impulse control may fundamentally shift over time.

The nature of this shift may relate to which actions and computations the animal must concentrate on in order to maximize reward. Initially, the rat must learn which options deliver the greatest yield through trial-and-error exploration. Throughout this phase, dopamine signals relevant to contingency tracking, reward prediction, and credit assignment should therefore dominate. Once the favoured choice strategy has been established, the animal is ready to exploit what it has learned. Now, profit maximisation depends on suppressing inappropriate or counter-productive responses, including those made prematurely at the nosepoke array. The animal must infer when to switch between these strategies, as the task structure and reinforcement contingencies remain constant throughout training.

Cholinergic interneurons within the striatum may play a key role in implementing such latent shifts in task-state. Their unique morphology and widespread connectivity make them ideally suited to this function. Although comprising only 2-5% of striatal neurons (Descarries *et al*., 1997; Mechawar *et al*., 2000), the axons and dendrites of CINs bifurcate extensively, such that the dendritic tree of a single neuron can span 500ums (Phelps *et al*., 1985; Wilson *et al*., 1990). Each CIN is estimated to produce around 500,000 axonal varicosities (Bolam *et al*., 1984; Contant *et al*., 1996), and the resulting axo-axonal signaling looks remarkably synapse-like (Kramer *et al*., 2022; Nosaka & Wickens, 2022). Indeed, CINs within the dorsomedial striatum encode latent task-states and facilitate the switching between behavioural strategies when unsignalled shifts in task-state occur (Stalnaker *et al*., 2016). Furthermore, it is now well-established that stimulation of CINs is sufficient to trigger dopamine release from terminal boutons within the dorsal and ventral striatum, independent of firing activity at dopaminergic cell bodies (Cachope *et al*., 2012; Threlfell *et al*., 2012). CINs can therefore directly gate the impact of striatal dopamine signaling in order to prioritize signals relevant for the present goals of the individual.

We therefore hypothesized that altering aCIN activity during task acquisition versus performance would predominantly affect decision-making versus motor impulsivity, consistent with our predictions of what may be the most important dopaminergic signal to prioritize in order to optimize behavior. We used DREADDs in combination with a ChAT-Cre line of rats to gain bidirectional control over aCINs in the NAC. In separate cohorts of female and male rats, we assessed either the effect of 1) chronic CIN modulation during learning of the cued rGT or 2) acute aCIN modulation following the establishment of stable decision making profiles.

## Methods

### Animals

All rats used for Experiment 1 (n = 88; n_female_ = 43) and Experiment 2 (n = 64; n_female_ = 33) were bred in-house from wildtype female Long-Evans (Charles River, St. Constant, QC) and transgenic ChAT::Cre bacterial artificial chromosome rats (LE-Tg(Chat-Cre)5.1Deis, RRRC, USA). Only those rats testing hemizygous for co-expression of Cre and choline acetyltransferase (ChAT) were used as control and experimental animals. Genotyping procedures and housing conditions are detailed in the supplemental material.

### Viral infusion surgery

At PND 35, experimental rats received bilateral infusions (1 μl / hemisphere) into the nucleus accumbens (NAc; AP: 1.7, ML: +/- 2.0, DV: −7.1 from skull) of AAV5-hSyn-DIO-hM3D(Gq)-mCherry or AAV5-hSyn-DIO-hM4D(Gi)-mCherry (7×10^12^ vg/mL; Addgene). Control rats received AAV5-hSyn-DIO-mCherry (7×10^12^ vg/mL; Addgene) (Figure 1A). Further surgical particulars are described in (Hynes *et al*., 2021).

**Figure 1.**
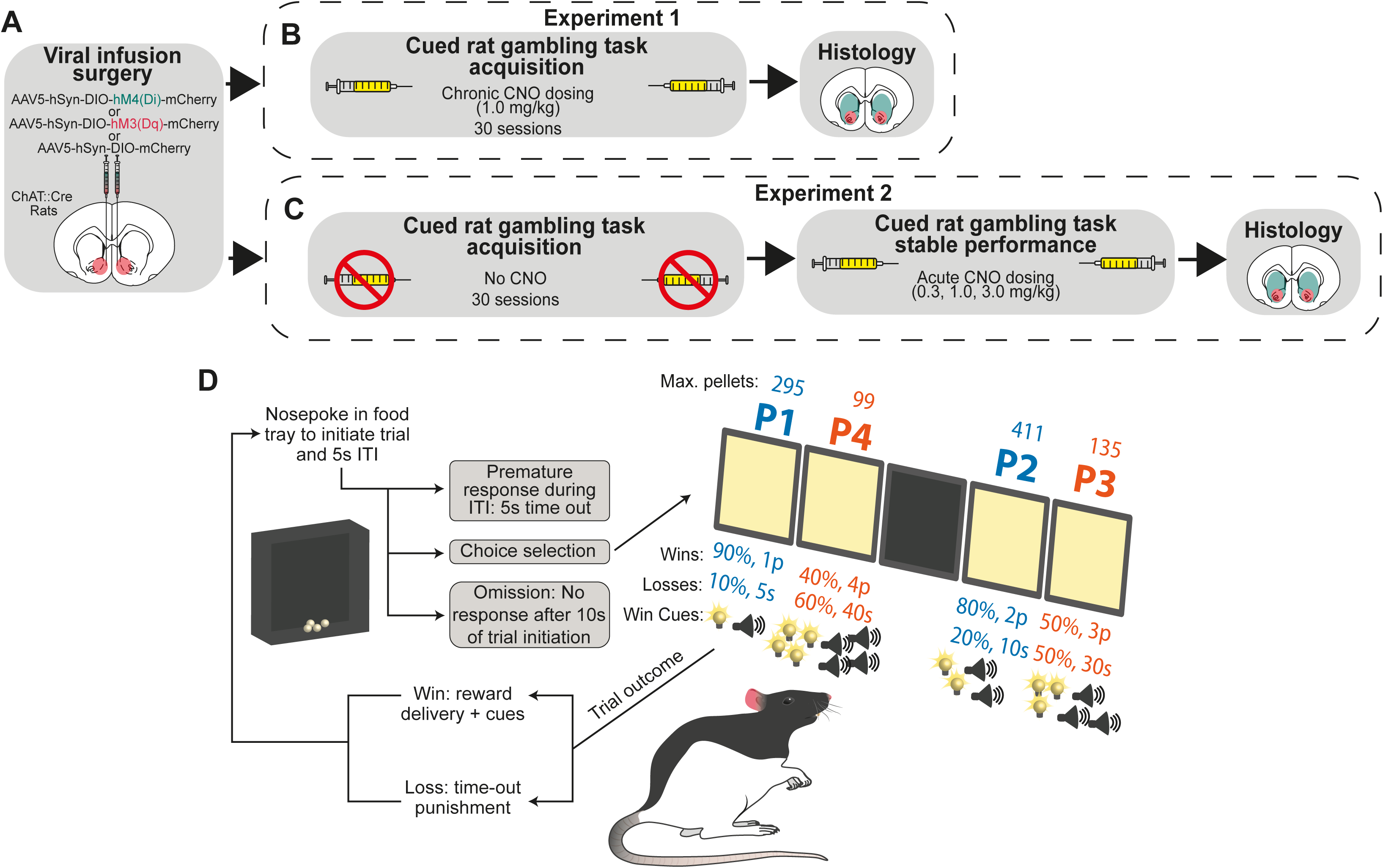
Experimental timeline and schematic of the cued rat gambling task (crGT). (A) Either an inhibitory or excitatory DREADD or a control fluorophore viral construct were delivered into the nucleus accumbens of female and male rats. After 4-6 weeks of recovery, (B) one group of rats were trained in the crGT under the daily influence of CNO (Experiment 1) and (C) another group was trained to stable performance in the crGT without administration of CNO (Experiment 2). Rats from experiment 2 underwent an acute CNO challenge. After the final crGT session, all rats were euthanized and expression of the viral construct was confirmed. (E) Schematic of the crGT task structure, which is described in detail in the methods section.

### Drugs

Desiccated CNO powder (Sigma Alrich) was dissolved in 100% dimethylsulfoxide (DMSO) and vortexed at room temperature until completely dissolved. The solution was then diluted with sterile saline to achieve a final vehicle concentration of 5% DMSO.

### Cued rat gambling task (crGT)

Full details of the behavioural apparatus and crGT pre-training procedure and conditions are outlined in the supplemental materials.

*Free choice training (30 sessions; **Experiment 1:** 1.0 mg/kg i.p. CNO 30 minutes before session start; **Experiment 2:** No CNO)*: During each 30-minute crGT session, rats sampled between four response holes, each of which was associated with distinct magnitudes and probabilities of sucrose pellet rewards or time-out punishments (see Figure 1 for specifications of probabilistic rewards and punishment). The optimal approach was to favour options which delivered smaller per-trial gains but lower time-out penalties, with the two-pellet choice (P2) resulting in the most reward earned per unit time. Overall decision making score was calculated as the difference between the summative probability of making a good choice and that of making a bad choice [i.e., score = (P1+P2) – (P3+P4)]. A 2s audiovisual cue was presented concurrently with reward delivery on each option which increased in salience and complexity with the size of the reward. The precise qualities of the stimuli are described in the supplemental materials. Responses made during the ITI were recorded as premature responses, a measure of impulsive action, which resulted in the illumination of the house light and a 5s time-out penalty after which a new trial could be initiated. If a response was not made into one of the 4 holes during the 10s stimulus presentation, the trial would be registered as an omission, and the tray light would be illuminated to signal another trial could be initiated. Each session lasted 30 minutes. More details on the crGT can be found in Barrus & Winstanley, 2017 and in (Figure 1B-D).

*Acute CNO challenge (**Experiment 2** only):* Once a stable, asymptotic behavioural baseline had been established over the last 5 sessions of CNO-free crGT training (achieved in 30 sessions), rats received either a vehicle control (5% DMSO in saline; VEH) or one of three different doses of CNO (0.3, 1.0, and 3.0 mg/kg, i.p.) according to a diagram balanced Latin Square design (Zeeb et al., 2009) (Figure 1C).

### Immunohistochemistry

To visualize DREADD expression in accumbal cholinergic interneurons, animals were anaesthetised and transcardially perfused with saline followed by 4% paraformaldehyde. 35-μm coronal striatal sections were then visualized for co-expression of mCherry and choline acetyltransferase (Figure 2A-F). The extent of mCherry transduction for each animal was then illustrated to scale on a striatal diagram adopted from Paxinos & Watson, 2009 (Figure 2G). Detailed information regarding the antibodies, incubation, and microscopy can be found in the supplemental materials.

**Figure 2.**
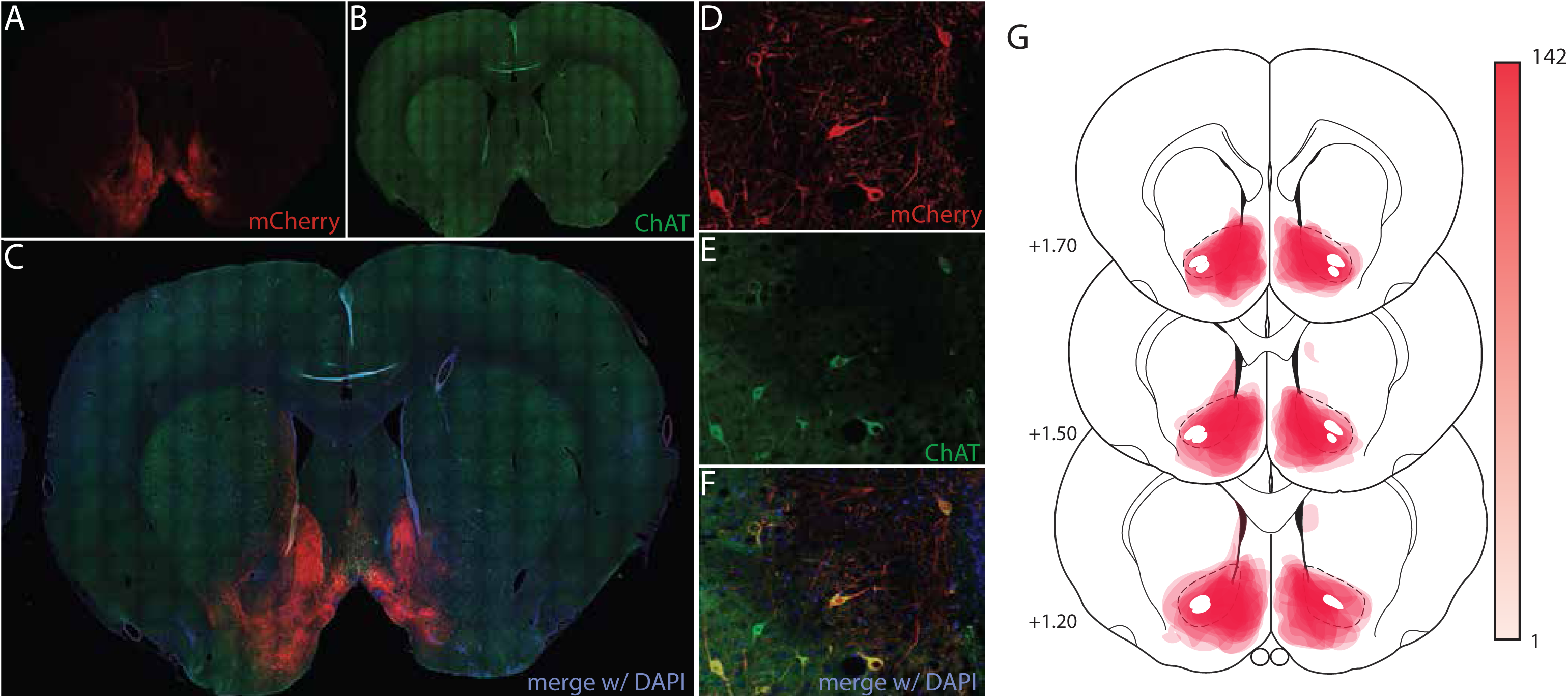
Selective expression of the viral constructs in cholinergic interneurons (CINs) of the nucleus accumbens (NAc). (A-C) Representative 10X tilescans of the entire striatum at AP +1.5 showing mCherry+ cells localised to the NAc. (D-F) 63X micrographs demonstrating viral transduction expression solely in CINs. (G) Illustrative reproduction of the spread of viral expression across all animals. Scale bar shows the relationship between colour intensity and the number of animals expressing mCherry in a specific location

### Statistical Analyses

For all dependent variables, two-way repeated measures ANOVAs with session or dose as the within-subjects factor and group (2 levels: experimental, control) as the between-subjects factor were used to detect omnibus effects. Significant terms in these ANOVAs were further probed using Fisher’s LSD post hoc comparisons to detect between-subjects differences at a given session in the chronic experiments, or differences from vehicle for a given dose in the acute experiments. Simple linear regression was also used to determine whether the slopes of the dependent variable differed across training between groups in Experiment 1.

## Results

### Immunohistochemistry

Expression of the viral constructs was largely limited to the NAC. Bilateral expression of mCherry was detected in the NAC core of all animals, with a substantial number also showing at least some expression in the shell subregion (Figure 2A-C). Only ChAT+ neurons expressed mCherry, suggesting selective viral transduction of cholinergic interneurons (Figure 2D-F). In a small minority of animals, a few mCherry+ cells were detected in the injector tract in the dorsal striatum (Figure 2G)

### Experiment 1 – Chronic modulation of aCINs during crGT acquisition

#### Decision making

Omnibus ANOVAs justified separate analysis of females and males for decision making score (session x sex x group: F_58,2088_ = 2.31, p < 0.0001) and for each individual choice (session x sex x group x choice: F_174,6264_ = 1.21, p = 0.032).

##### Females

All females underwent the longitudinal decline in decision making score that is normally observed in the crGT (Hynes et al., 2021; Hynes et al., 2024; Mortazavi et al., 2024) (all slopes < 0; test of non-zero slopes: all p < 0.009), but the rate of decline was increased by chronic aCIN inhibition (slope_hM4_ vs. slope_control_: F_2,1284_ = 10.53, p < 0.001), resulting in a decision making score that was significantly lower than controls (session x group: F_58,1160_ = 2.70, p < 0.001) from session 8 onward (hM4_s1-s7_ vs control_s1-s7_: all ps ≥ 0.130; hM4_s8-s30_ vs control_s8-s30_: all ps ≤ 0.047) (Figure 3A). This decrease in score was driven by a reduction in P2 choice (session x group: F_58,1160_ = 2.13, p < 0.0001) from session 8 onward (hM4_s1-s7_ vs control_s1-s7_: all ps ≥ 0.144; hM4_s8-s30_ vs control_s8-s30_: all ps ≤ 0.050) (Figure 3B) and a linear increase in P4 (Figure 3E) (session x group: F_58,1160_ = 2.13, p < 0.0001; slope_hM4_ vs. slope_control_: F_1,896_ = 6.51, p = 0.011). Choice of P1 (session x group: F_58,1160_ = 0.88, p = 0.726) and P3 (session x group: F_58,1160_ = 0.90, p = 0.691) did not differ between groups (Figure 3B,D).

**Figure 3.**
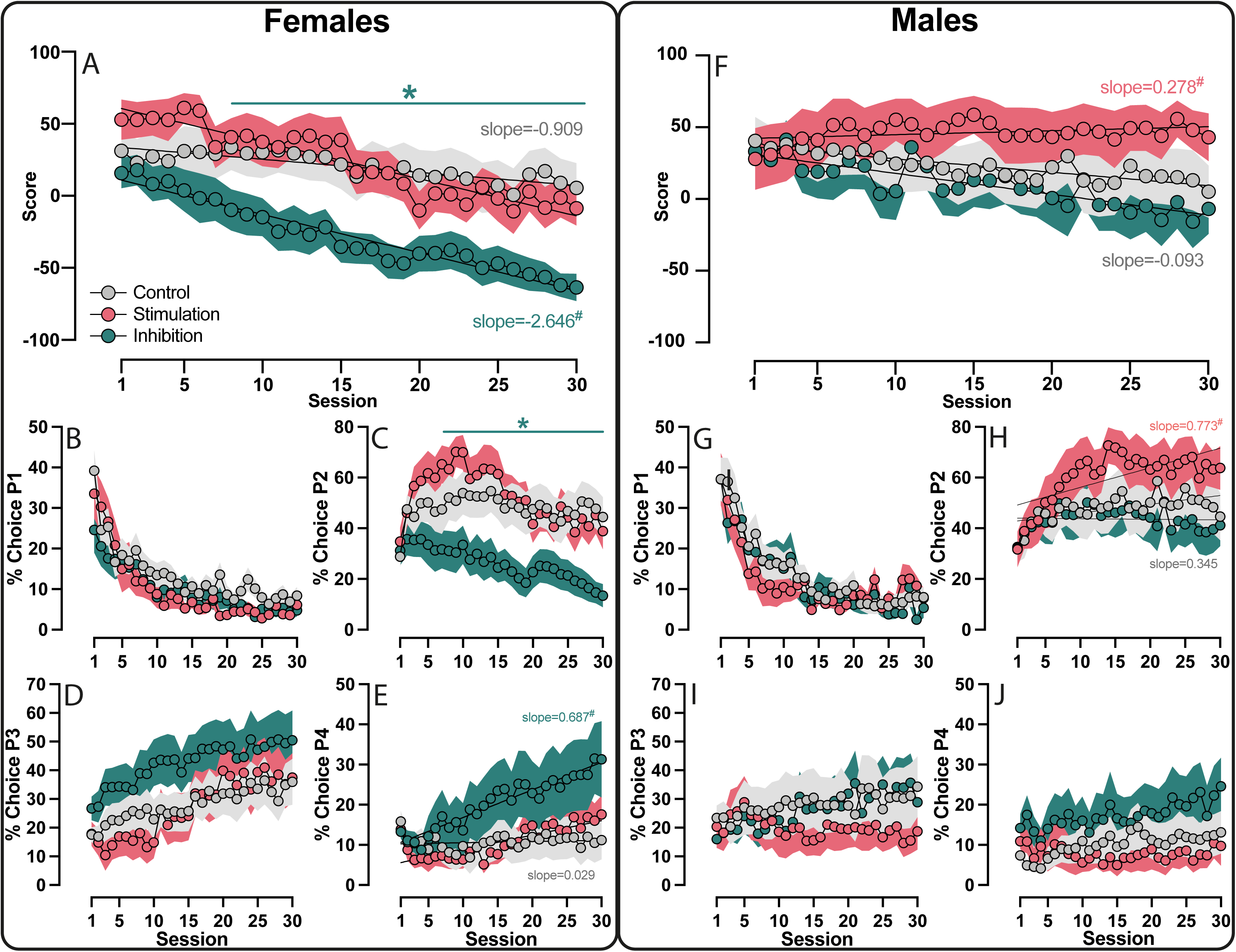
Chronic modulation of accumbal cholinergic interneurons (aCINs) has sex-dependent effects on the development of a decision making profile. (A) Inhibition of aCINs exacerbates the normal longitudinal decline in decision making score in females, (C) by reducing the optimal choice of P2 and (E) increasing the risky choice P4. (B) P1 and (D) P3 were unaffected by any manipulation. (C) Stimulation of aCINs transiently increased the preference for P2 in females. (F) In males, stimulation of aCINs prevents the longitudinal decline in decision making score by (H) increasing the preference for P2. (F-G) Inhibition of aCINs has no effect on decision making in males. # p < 0.05 between-subjects difference in slope difference versus control. * indicates p < 0.05 between-subjects difference in median score versus control. # indicates that the slope is significantly different from controls

Chronic stimulation of aCINs in females did not influence the overall decision making score but did cause a transient increase in choice of P2 in the first 10 sessions (slope_hM4(s1-s10)_ vs. slope_control(s1-s10)_: F_1,56_ = 58.29, p < 0.001) (Figure 3C).

##### Males

Chronic stimulation of aCINs prevented the expected longitudinal decline in decision making score observed in control rats and those expressing the inhibitory DREADD (difference of slopes: F_2,1044_ = 5.81, p = 0.003; slope_control_ = −0.934, slope_HM4_ = −1.454, slope_HM3_ = 0.278) (Figure 3F). Underlying this observation was an increased preference for P2 over time in stimulated rats (slope_HM3_ vs. slope_control_: F_1,687_ = 3.45, p = 0.011) (Figure 3H). Choice of P1 (session x group: F_58,928_ = 0.877, p = 0.877), P3 (session x group: F_58,928_ = 1.21, p = 0.4384), and P4 (session x group: F_58,928_ = 0.717, p = 0.9453) did not differ between groups (Figure 3G, I & J).

#### Impulsivity

As expected, based on our previous reports(Hynes *et al*., 2021; Hynes *et al*., 2024b) and justifying the *a priori* independent analysis of either sex, control females made fewer premature responses than their male counterparts (sex_control_: F_1,27_ = 8.31, p = 0.0080) (Figure S1A & S1B). Chronic modulation of aCIN activity during task acquisition did not alter premature responding in neither females (group: F_2,40_ = 0.984, p = 0.383) nor males (group: F_2,32_ = 2.091, p = 0.140).

#### Other Variables

See supplemental materials.

### Experiment 2 – Acute modulation of accumbal CINs following crGT acquisition

#### Decision making

In stark contrast to the results of Experiment 1, aCIN modulation after crGT performance was well-established had no effect on decision score in either sex (Figure 4A & B; session x group x dose-females: F_6,90_ = 1.68, p = 0.1359; - males: F_6,84_ = 0.33, p = 0.9207; Figures 4A & 4B). In line with these results, aCIN modulation also did not alter preference for any specific option (session x group x choice x dose-females: F_18,270_ = 1.68, p = 0.1359;-males: F_18,252_ = 0.59, p = 0.9060).

**Figure 4.**
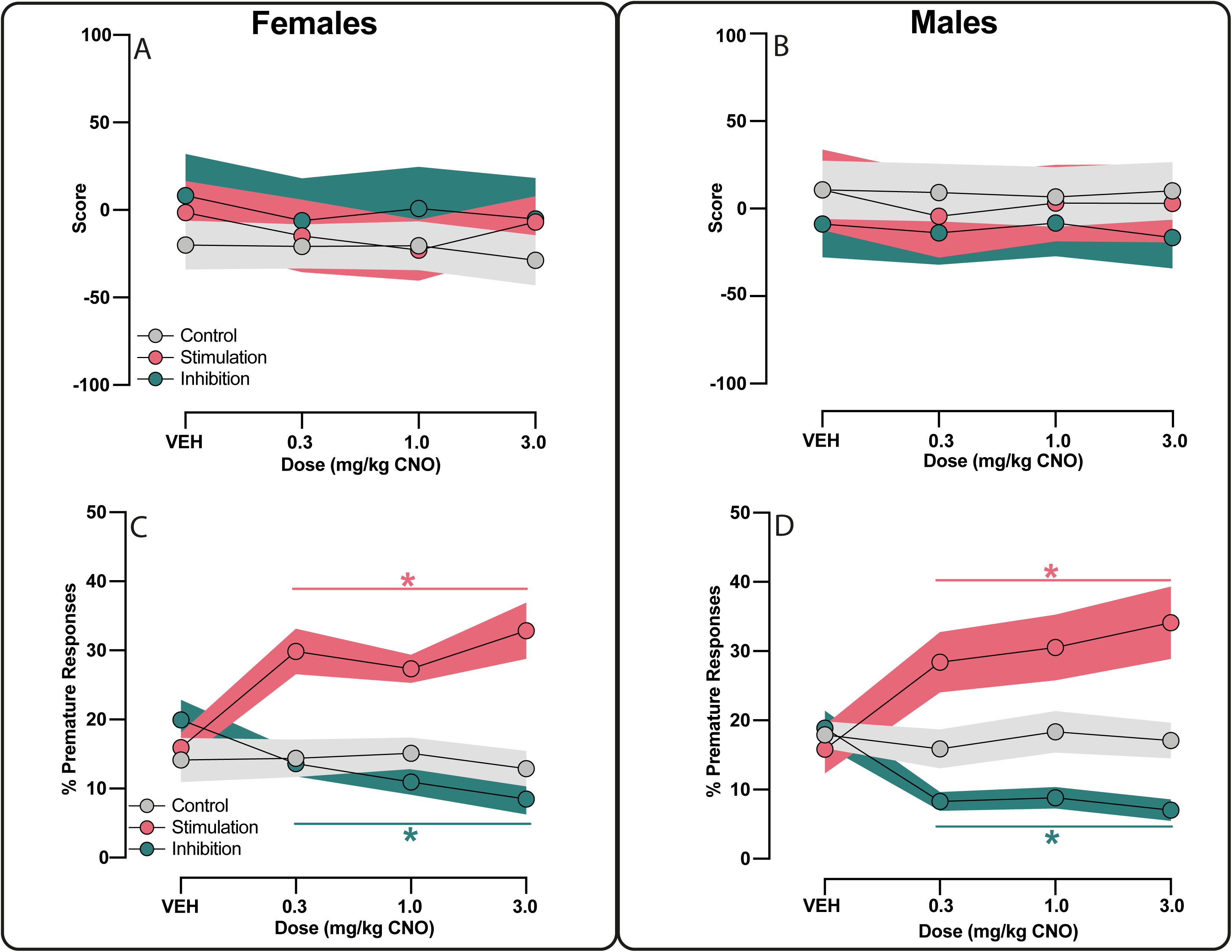
Chronic modulation of accumbal cholinergic interneurons (aCINs) does not affect decision making but modulates motor impulsivity similarly in females and males. Once stable crGT performance was established, in neither (A) females nor (B) males did modulation of aCINs affect decision making score. Stimulation of aCINs increased premature responding at all doses, whereas inhibition of aCINs had the opposite effect, in both (C) females and (D) males. “*” indicates within-subjects p < 0.02 compared to VEH.

#### Impulsivity

Whereas aCIN manipulation during task acquisition did not systematically alter premature response rates in either sex, once rats had learned the task, stimulation of aCINs increased premature responding at all doses in both females and males (females: dose x group: F_6,90_ = 14.34, p < 0.001; 0.3 vs VEH: p = 0.004; 1.0 vs VEH: p = 0.002; 3.0 vs VEH: p < 0.001; males: dose x group: F_6,78_ = 13.38, p < 0.001; 0.3 vs VEH: p = 0.009; 1.0 vs VEH: p = 0.003; 3.0 vs VEH: p = 0.002). Conversely, inhibition of aCINs decreased premature responding in both sexes (females: 0.3 vs VEH: p = 0.023; 1.0 vs VEH: p = 0.004; 3.0 vs VEH: p < 0.001; males: 0.3 vs VEH: p = 0.002; 1.0 vs VEH: p = 0.002; 3.0 vs VEH: p < 0.001). CNO did not affect premature responding in control females (Figure 4C; 0.3 vs VEH: p = 0.874; 1.0 vs VEH: p = 0.589; 3.0 vs VEH: p = 0.251) or control males (Figure 4D: 0.3 vs VEH: p = 0.152; 1.0 vs VEH: p = 0.820; 3.0 vs VEH: p < 0.525).

#### Other variables

See supplemental materials.

## Discussion

Here we show that bi-directional modulation of aCINs significantly affected either motor impulsivity or decision making, depending on whether aCINs were targeted after a stable asymptotic baseline had been established on the crGT, or while behaviour was still in flux during task acquisition. These results point to divergent effects of cholinergic modulation of accumbal function across different task states, allowing signals relevant for current goals to dominate. Unpacking this double dissociation further, we found that chemogenetic stimulation of aCINs increased motor impulsivity, while chemogenetic inhibition suppressed this behaviour, when the manipulation was delivered during stable task performance. This effect was not accompanied by shifts in choice patterns, and was observed in animals of both sexes. In contrast, manipulating aCIN activity while animals were learning the crGT did not affect motor impulsivity, but instead shifted choice preferences in a sex-dependent manner. Specifically, chemogenetic inhibition of aCINs amplified risky choice in female rats, while chemogenetic activation had no effect. Conversely, in male rats, chemogenetic inhibition of aCINs had no effect on the evolution of choice patterns, while chemogenetic activation reduced risky choice.

Our observation that modulating aCINs affected decision making during acquisition, but impulsivity once choice strategies had been learned, is in keeping with our original hypothesis that aCINs may gate dopamine signalling according to latent task state, but what could underlie this shift? The locus of dopaminergic control over behaviour shifts from the ventral to dorsal striatum as a task becomes well-learned and switches from goal-directed to habitual (Atallah *et al*., 2007; Graybiel, 2008; Smith & Graybiel, 2016). It may be inferred behaviourally that this process also occurs in the cued rGT, as decision-making behaviour becomes insensitive to reinforcer devaluation after extensive training, suggesting a loss of goal-directed control (Hathaway *et al*., 2021). As such, aCIN modulation of dopaminergic signals may have failed to alter decision making once the task had been well-learned, because the locus of control of decision making had shifted dorsally away from the NAc. In the dorsomedial striatum, frontal input to CINs can influence the behavioural response to latent shifts in task state (Stalnaker *et al*., 2016). However, whether frontal input to CINs in the NAc is critical for the shift from goal-directed to habitual control of behaviour has yet to be determined. The current results do not tell us whether aCINs have a causal role in determining which dopaminergic signals are important for behaviour, or simply modulate existing signals prioritised through an alternate mechanism.

Females and males reacted similarly to CIN activation and inhibition after the task had been learned, indicating that CINs do not necessarily function in a fundamentally different manner across the sexes. We have previously observed dramatic sex differences in the response to chronic chemogenetic activation of dopaminergic neurons of the VTA during acquisition of the cued rGT; females developed a risky strategy more slowly, whereas this was evident more rapidly in males (Hynes *et al*., 2024b). However, in the absence of win-paired cues, all rats made significantly more risky choices throughout learning. As such, the presence of reward-concurrent audiovisual stimuli may be unmasking sex differences in the dopaminergic regulation of decision-making processes within the mesolimbic pathway. The ability of CINs to regulate the balance of tonic-to-phasic dopamine signals by modulating the release of dopamine at terminal boutons (Cachope & Cheer, 2014) could be critical in explaining this pattern of effects.

Under normal conditions, aCINs are tonically active and maintain constant levels of local acetylcholine (ACh), which agonizes B2-subunit containing nicotinic acetylcholine receptors (nAChR_B2_). This causes depolarizing currents in dopamine terminals and the consequent continual release of DA (Morris *et al*., 2004). Activation and inhibition of aCINs can increase and decrease tonic accumbal DA levels, respectively, via this mechanism (Cachope & Cheer, 2014). Acute systemic and local accumbal pharmacological potentiation of dopamine neurotransmission increases premature responding in the 5CSRTT (Economidou *et al*., 2012; Moreno *et al*., 2013; Fitzpatrick *et al*., 2018). In contrast, inhibition of accumbal dopamine activity, through either pharmacological or chemogenetic methods, can decrease premature responding (Cole & Robbins, 1989; Hynes *et al*., 2020). If we assume that aCINs are primarily influencing premature responding by altering tonic levels of dopamine release, stimulation of aCINs should increase motor impulsivity and inhibition of aCINs should decrease motor impulsivity, just as we observe in the present acute manipulations. It is moreover unsurprising that we observed no sex differences in the acute effects on impulsivity, as females and males do not appear to differ in baseline levels of tonic dopamine (Rivera-Garcia *et al*., 2020; Kuiper *et al*., 2024). There are, however, a substantial number of reports of sex differences in phasic and evoked striatal dopamine release (Walker *et al*., 1999; Dluzen & McDermott, 2008; Zachry *et al*., 2021).

In the presence of motivationally relevant stimuli, such as reward-paired cues, aCINs pause firing and dopamine neurons coincidentally burst fire. In concert, this increases the so-called “signal-to-noise” of phasic dopamine and amplifies its efficacy as a teaching signal, or more generally, an effector of physiological adaptation in the striatum (Franklin & Frank, 2015; Nougaret & Ravel, 2015). As such, chronic inhibition of aCINs should enhance the physiological impact of phasic dopamine in the striatum by reducing tonic dopamine levels, whereas stimulation of aCINs should blunt the effect of phasic dopamine signalling by increasing dopaminergic tone. Both chemogenetic and optogenetic stimulation of aCIN activity suppresses the expression of Pavlovian-to-instrumental transfer through the activity of ACh at DA terminal nAChRs, suggesting that aCINs oppose the motivational effect of cues by augmenting terminal dopamine release (Collins *et al*., 2019). We have previously theorised that the relatively greater levels of risky choice seen in the cued vs uncued version of the rGT results from cue-evoked phasic dopaminergic events (Winstanley & Hynes, 2021). Inhibiting aCINs should therefore amplify the influence of these phasic signals and potentiate risky choice whereas stimulation of aCINs should blunt the physiological impact of phasic dopamine signalling in the accumbens, and attenuate risky decision making. These predictions are confirmed in our results, yet the question remains as to why the former is observed only in females and the latter exclusively in males.

Females display enhanced electrically-evoked phasic dopamine transients, which are subject to greater reuptake rates (Walker *et al*., 1999), resulting in a phasic dopamine signal that is of greater amplitude and narrower width compared to males. As depicted in Figure 5, decreasing terminal dopamine release through inhibition of aCINs may therefore have been sufficient to amplify risky choice in females, as the phasic signals are more finely tuned, while increasing terminal dopamine release through stimulation of aCINs would have minimal effects. Conversely, the broader phasic dopamine peaks in males may be less affected by subtle decreases in tonic dopamine levels caused by aCIN inhibition, but more easily masked by greater terminal dopamine release caused by aCIN stimulation. Although speculative, this theory provides a plausible explanation for the pattern of results reported here that can be tested in future experiments.

**Figure 5.**
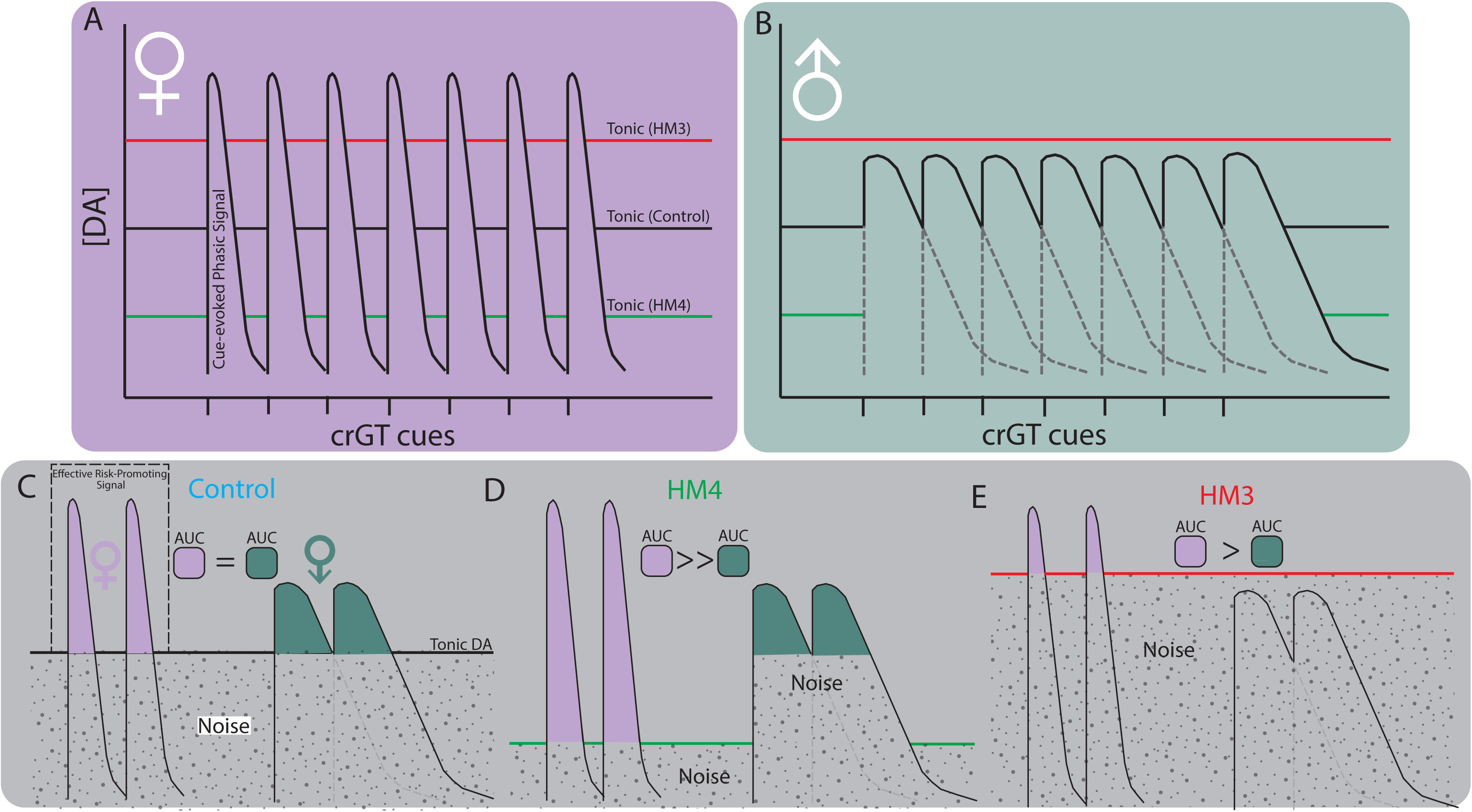
Accumbal cholinergic interneuron (aCIN) modulation of tonic dopamine differentially affects the signal-to-noise ratio of the cue-evoked risk-promoting phasic dopamine signal in a sex-dependent manner. (A) Females and (B) males do not differ in baseline tonic dopamine (DA) levels nor in the degree to which chemogenetic modulation of aCINs affects tonic DA levels. The profile of cue-evoked phasic DA transients differs substantially between the sexes (Walker *et al*., 1999), with females having higher amplitude and more rapidly assimilated transients than males. The green line illustrates that stimulation of aCINs increases tonic DA levels and the red line shows that inhibition of aCINs decreases tonic DA. (C) When aCINs are unmanipulated, the area under the curve (AUC) corresponding to the effective risk-promoting phasic dopaminergic signal does not differ between females and males, so both sexes display similar a cue-induced longitudinal decline in decision making score. (D) When the tonic DA level is decreased via aCIN inhibition, the cue-evoked phasic DA signal stands out amid the reduced tonic noise, but the effective risk promoting signal is greater in females, because slower DA reuptake in males prevents the complete resolution of temporally proximal transients. Thus, only in females does aCIN inhibition promote risk-taking. (E) Increasing the tonic DA levels completely masks the phasic signal in males, thereby preventing cue-induced decision making deficits. The phasic signal in females is substantially reduced but still evident, resulting in a transient and moderate attenuation of cue-induced risk-taking.

Our present focus on pre-synaptic mechanisms of acetylcholine-evoked terminal dopamine release may seem to overlook the possibility that acetylcholine could directly influence the output projection circuity of the striatum via post-synaptic receptors. Indeed, both direct and indirect pathway-projecting accumbal medium spiny neurons (MSNs) express muscarinic acetylcholine receptors (mAChRs)(Hersch *et al*., 1994; Ince *et al*., 1997), suggesting that their activity could be modulated by aCIN-derived acetylcholine. However, striatal MSN activity is not affected by muscarine or the broad mAChR antagonist scopolamine in dopamine-depleted striatal slices (Dehorter *et al*., 2009), suggesting that dopamine is necessary for a physiologically relevant effect of acetylcholine on striatal MSNs.

A more likely co-contributor to the behavioural effects we report here are striatal astrocytes. Though hitherto understudied, striatal astrocytes are emerging as important regulators of motivated behaviour. Each striatal astrocyte interacts with thousands of striatal dopaminergic synapses, expresses nAChRs (Delbro *et al*., 2009; Grybko *et al*., 2010; Adermark & Bowers, 2016) and exhibits robust changes in calcium and potassium signalling following exposure to nicotine (Oikawa *et al*., 2005; Hernández-Morales & García-Colunga, 2009). As such, astrocytes are well-positioned to fine-tune dopamine release (Corkrum & Araque, 2021) and reuptake (Hynes *et al*., 2024a) in response to cholinergic modulation. Considering that sex differences in evoked dopamine are largely attributable to sex differences in the expression of DAT (Zachry *et al*., 2021), and that cue-controlled behaviour has recently been discovered to be associated with the expression of astrocytic DAT (Hynes *et al*., 2024a; Fouyssac *et al*., 2025), the behavioural processes driven by striatal ACh-DA interactions may be, at least in part, governed by the local astrocytic syncytium. Indeed, it has been elsewhere suggested that striatal astrocytic mechanisms may influence decision making in tasks such as the present one (Hynes *et al*., 2025).

There are two important caveats to the interpretation of the results we have presented here. Firstly, our viral manipulation was not restricted to the core or shell region of the NAc. Given that the NAcC codes the instantaneous incentive value of reward cues, while the NAcS detects emerging changes in the incentive value of cues (Floresco *et al*., 2008; Ambroggi *et al*., 2011), the NAcS may be of minor relevance in the crGT, as the cue-reward contingencies never change crGT. As such, the observations we report here are likely to be predominantly NAcC-mediated.

The second caveat is that we do not account for the effect of circulating gonadal hormones. Though hormonal load (as inferred from the oestrous stage) does not impact decision making and nor do females display overall differences or greater longitudinal variability in decision making compared to males (Hynes *et al*., 2020; Hynes *et al*., 2021; Hynes *et al*., 2024b), heightened motor impulsivity is observed during the proestrus phase (Hynes *et al*., 2020), so hormonal factors may play into the outcome our acute manipulations. To fully disambiguate the hormonal versus organizational sex differences in the aCIN control of impulsivity, similar acute experiments would need to be carried out in gonadectomized rats.

In summary, we present evidence that, depending on task state, aCINs subserve dissociable roles in impulsivity and decision making, the latter of which differs across the sexes. We furthermore put forth a testable set of predictions that combines the established neurobiology of accumbal ACh-DA interactions with our emerging hypotheses of how reward cues impair decision making. These predictions aim to reconcile a) why decision making and impulsivity are differentially affected by the time course of aCIN modulation and b) why sex differences are observed only in the decision making facet of aCIN-modulated behaviour. Of course, these predictions remain to be substantiated with *in vivo* monitoring of accumbal ACh-DA dynamics in behaving animals, but may serve to reveal novel therapeutic targets for the treatment of addiction, ADHD, and other psychiatric conditions in the impulsive-compulsive domain.

## Author contributions

TH and CW designed the experiments. TH, CC, KH, GB, AL, and LC carried out the behavioural experiments. TH and CC conducted the histology. TH, CC, and CW analyzed and interpreted the data and wrote the manuscript.

## Acknowledgements and funding

The experimental work was carried out in The Department of Psychology and The Djavad Mowafaghian Centre for Brain Health at The University of British Columbia (located on the unceded ancestral lands of the Coast Salish people) and was funded by a CIHR grant to CW (PJ-162312), an Institute for Mental Health Marshall Doctoral Scholarship to TH and a CGS-M scholarship to CC. TH and CC carried out the writing of the manuscript at The University of Cambridge, where TH is funded by a Leverhulme Early Career Fellowship (ECF-2024-242) and an Isaac Newton Trust Fellowship (24.08). CC is funded by a Cambridge Trust International Scholarship (10712081 – 2023) and an NSERC PGS-D (PGSD3 - 579465 – 2023).

