## Supplemental Materials for "Accumbal cholinergic interneurons regulate decision making or motor impulsivity depending on latent task state"

**Supplemental Methods**

*Animals – genotyping and housing conditions.* Expression of Cre-recombinase was confirmed via PCR. To extract DNA, ear notch samples were lysed using a buffer solution (50 mM pH 8.0 Tris, 2 mM NaCl,10 mM EDTA, 1% SDS) and Proteinase K (Invitrogen, part number 25530-015). Extracted DNA was stored at -20°C until PCR was performed. All reactions were performed in 200 μl thin-walled PCR tubes with FASTSTART 28 TAQ DNA POL. DNTPACK reaction mix (Roche, cat#4738357001), primers for Cre (forward: 5’-AGA GTA CAC TGT GGG CAG GA-3’; reverse:5’-GCA AAC GGA CAG AAG CAT TT- 3’) or ChAT (forward: 5’-GTG GCT CAG AAC AGC AGC ATC A-3’; reverse: 5’-CCT CAC TGA GAC GGC GGA AAT T-3’). Samples were then placed in a thermocycler and underwent standard cycling protocols (95°C 5 mins; then cycled 35 times: 94°C 30 sec, 63°C 30 sec, 72°C 1 min; holding at 72°C for 10 min; infinite holding at 4°C) and subsequently run on an agarose gel at 90V for 40 min to verify genotypic expression. Rats were weaned at postnatal day 21 and pair- or trio-housed with same sex littermates in a climate-controlled colony room on a reverse 12-hour light-dark cycle (lights off 08:00; temperature 21°C). One week before the commencement of behavioural training, rats were food restricted to 85% of their theoretical free-feeding weight and continued to be food-restricted throughout the experiment.

*Behavioural apparatus.* The cued rat gambling task (crGT) behavioural assay was conducted in 32 operant conditioning chambers (30.5 × 24 × 21 cm; Med Associates, St. Albans, VT, USA), located in separate rooms. Set in the curved wall of each box was an array of five nose-poke holes. Each nose-poke unit was equipped with an infrared detector and a yellow light-emitting diode stimulus light. Chow pellets (45 mg, Formula P; Bio-Serv) could be delivered at the opposite wall via a dispenser. Online control of the apparatus and data collection was performed using code written by CAW in MEDPC (Med Associates) running on standard IBM-compatible computers. Each chamber was enclosed within a ventilated sound-attenuating cabinet (Med Associates Inc, VT) equipped with a fan to provide ventilation and further mask extraneous noise.

*Cued rat gambling task pre-training procedures.*

Habituation to the operant box (1 session; Experiments 1 & 2: no CNO): Animals were initially habituated to the operant chambers over one 30-minute exposure, during which sucrose pellets were placed in each of the apertures and animals were allowed to explore the apparatus.

Modified five-choice serial-reaction time task (~10 sessions; Experiments 1 & 2: no CNO): To encourage rats to nose poke in the illuminated array, animals were then trained on a variant of the five-choice serial-reaction time task (5-CSRTT). Animals initiated each trial by making a nosepoke response at the food magazine, which started the 5s intertrial interval (ITI). Following the ITI, one of the four response hole lights used in the crGT (described below) was pseudo-randomly illuminated for 10s. A nose-poke in the illuminated hole response was rewarded with a single sucrose pellet delivered to the food magazine. Incorrect responses, failure to respond at the illuminated aperture, or responding prematurely before the apertures were illuminated resulted in a 5s time-out punishment. Each session consisted of a maximum of 100 trials and lasted 30 min. Animals were trained on this task until responding reached 80% accuracy and 20% omissions.

Forced choice training (7 sessions; **Experiment 1**: 1.0 mg/kg i.p. CNO 30 minutes before session start; **Experiment 2:** No CNO): This training procedure ensured equal exposure to the different reinforcement contingencies associated with each aperture. This stage is identical to the crGT training described in the main text, except that only one aperture is illuminated on each trial. Roughly equal numbers of each trial type are presented in each session, thereby ensuring rats adequately sample all four options. Each session lasted 30 min.

Audiovisual stimuli qualities: P1 win: P1 hole flashes at 1 Hz, monotone; P2 win: P2 hole flashes for 1 Hz, tone changes pitch once after 1s; P3 win: P3 hole flashes at 5 Hz for 1s, followed by flashing of the two neighbouring holes in one of two patterns chosen at random, traylight flashes concurrently at 5 Hz for 2s, three different tones used, changing pitch every 0.1 s, in one of two patterns chosen at random; P4 win: P4 hole flashes at 5 Hz for 1s, followed by flashing of all five holes in one of four patterns at random, traylight flashes concurrently at 5 Hz for 2s, six different tones used, changing pitch every 0.1s, in one of four patterns chosen at random.

*Immunohistochemistry.* Coronal sections containing the striatum were incubated with primary antibodies against mCherry (Cat#ab205402; Abcam; Toronto, ON, Canada; 1:700 for 48h) and choline acetyltransferase (Cat#MA5-32663; Invitrogen; Thermo Fischer Scientific; Burnaby, BC, Canada; 1:250 for 48h) at 4°C. Sections were then washed in PBS and incubated with secondary antibodies conjugated to Alexa Fluor® 488 (Cat#A-21103) and Alexa Fluor® 633 (Cat#A-11034) (Thermo Fischer Scientific; Burnaby, BC, Canada; 1:1000 for both) for 2h at RT. Sections were then cover-slipped under HARLECO® KrystalonTM mounting medium (Thermo Fischer Scientific; Burnaby, BC, Canada) and visualized using an SP8 WLL confocal microscope (Leica Microsystems, Germany).

**Supplemental Results**

**Experiment 1 – Chronic modulation of aCINs during crGT acquisition**

Trials

*Females.* Modulation of accumbal CINs had no effect on the number of trials completed in females (session x group: F_58,1160_ = 1.073, p = 0.334; group: F_2,40_ = 1.108, p = 0.349) (Figure S1A).

*Males.* On session 1, HM3 males completed fewer trials than HM4 rats [F_58,928_ = 1.369, p = 0.038; p_HM3 vs. HM4_ = 0.046) (Figure S2B). On sessions 25 (p = 0.046) and 29 (p = 0.042), HM3 rats completed more trials than HM4 rats. We did not interpret these differences as meaningfully in the scope of this experiment.

Omissions

Modulation of accumbal CINs did not affect the number of trials omitted in females (session x group: F_58,1160_ = 1.130, p = 0.238; group: F_2,40_ = 0.312, p = 0.734) or males (session x group: F_58, 928_ = 1.478, p = 0.075; group: F_2,32_ = 3.065; p = 0.061) (Figures S2C & S2D).

Choice latency

Modulation of accumbal CINs did not affect choice latency in females (session x group: F_58,1160_ = 0.751, p = 0.916; group: F_2,40_ = 1.152, p = 0.326) or males (session x group: F_58, 928_ = 1.220, p = 0.131; group: F_2,32_ = 1.600; p = 0.218) (Figures S2E & S2F).

Collection latency

Modulation of accumbal CINs did not affect collection latency in females (session x group: F_58,1160_ = 1.104, p = 0.144; group: F_2,40_ = 0.482, p = 0.542) or males (session x group: F_58, 928_ = 0.959, p = 0.564; group: F_2,32_ = 2.688; p = 0.084) (Figures S2G & S2H).

**Experiment 2 – Acute modulation of accumbal CINs following crGT acquisition**

Trials

Acute modulation of accumbal CINs did not impact the number of trials completed by females (group x dose: F_6,90_ = 2.099, p = 0.691) or males (group x dose: F_6,84_ = 0.794, p = 0.577) (Figures S3A & S3B).

Omissions

Acute modulation of accumbal CINs did not impact the number of trials omitted by females (group x dose: F_6,90_ = 0.921, p = 0.484) or males (group x dose: F_6,87_ = 0.192, p = 0.978) (Figures S3C & S3D).

Choice latency

*Females*. Acute modulation of accumbal CINs did not affect choice latency (group x dose: F_6,90_ = 1.048, p = 0.400), but those expressing either chemogenetic construct were slightly faster than controls to make a choice (HM4 vs. Control: F_1,23_ = 15.06, p < 0.001; HM3 vs. Control: F_1,23_ = 3.982, p = 0.049) (Figure S3E).

*Males.* Acute modulation of accumbal CINs did not affect choice latency (group x dose: F_6,84_ = 1.916, p = 0.087) (Figure S3F).

Collect Latency

Acute modulation of accumbal CINs did not affect choice latency in females (group x dose: F_6,90_ = 1.434, p = 0.210) or males (group x dose: F_6,84_ = 0.491, p = 0.813) (Figures S3G & S3H).

**Figure S1.** *Chronic modulation of accumbal cholinergic interneurons (aCINs) does not affect impulsivity.* In neither (A) females nor (B) males did modulation of aCINs affect premature responding. Males are more impulsive than males.

**Figure S2.** *Effect of chronic aCIN modulation on other variables.* (A-B) Trials completed, (C-D) trials omitted, (E-F) latency to make a decision, (G-H) and latency to collect the reward.

**Figure S3.** *Effect of acute aCIN modulation on other variables.* (A-B) Trials completed, (C-D) trials omitted, (E-F) latency to make a decision, (G-H) and latency to collect the reward.
