## Supplementary figures and images for "Accumbal cholinergic interneurons regulate decision making or motor impulsivity depending on latent task state"

### Supplemental Figure 1

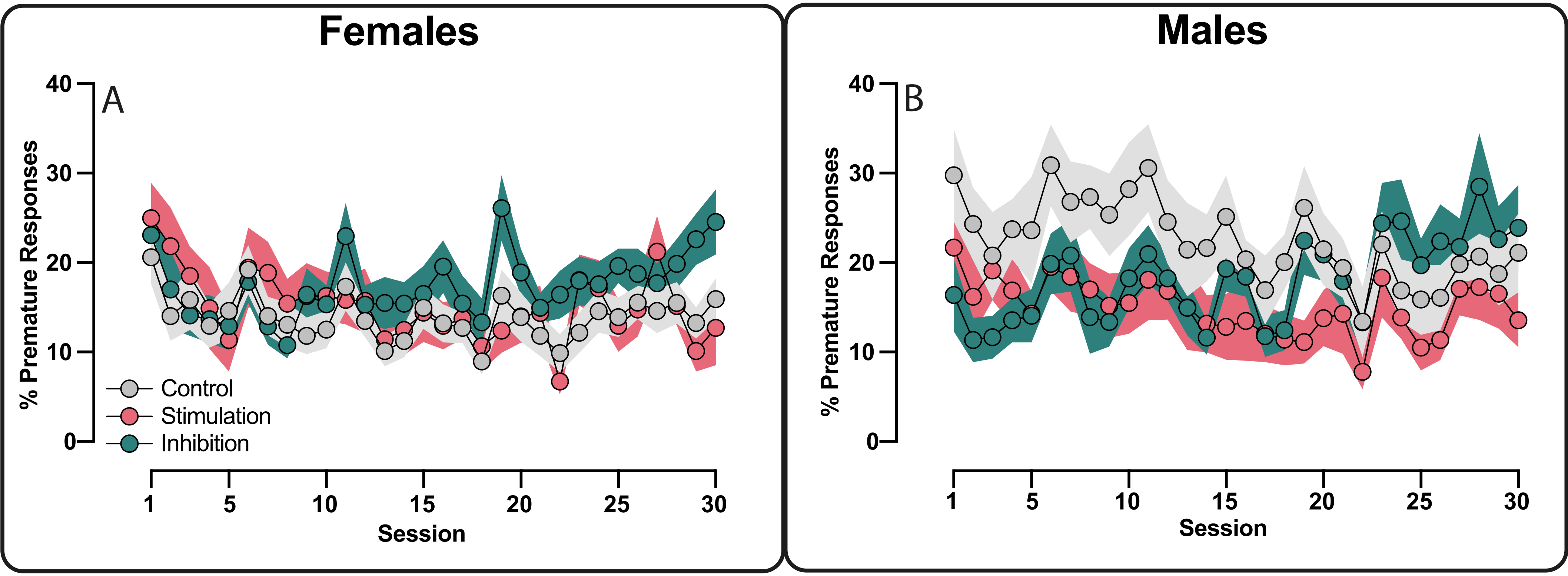

### Supplemental Figure 2

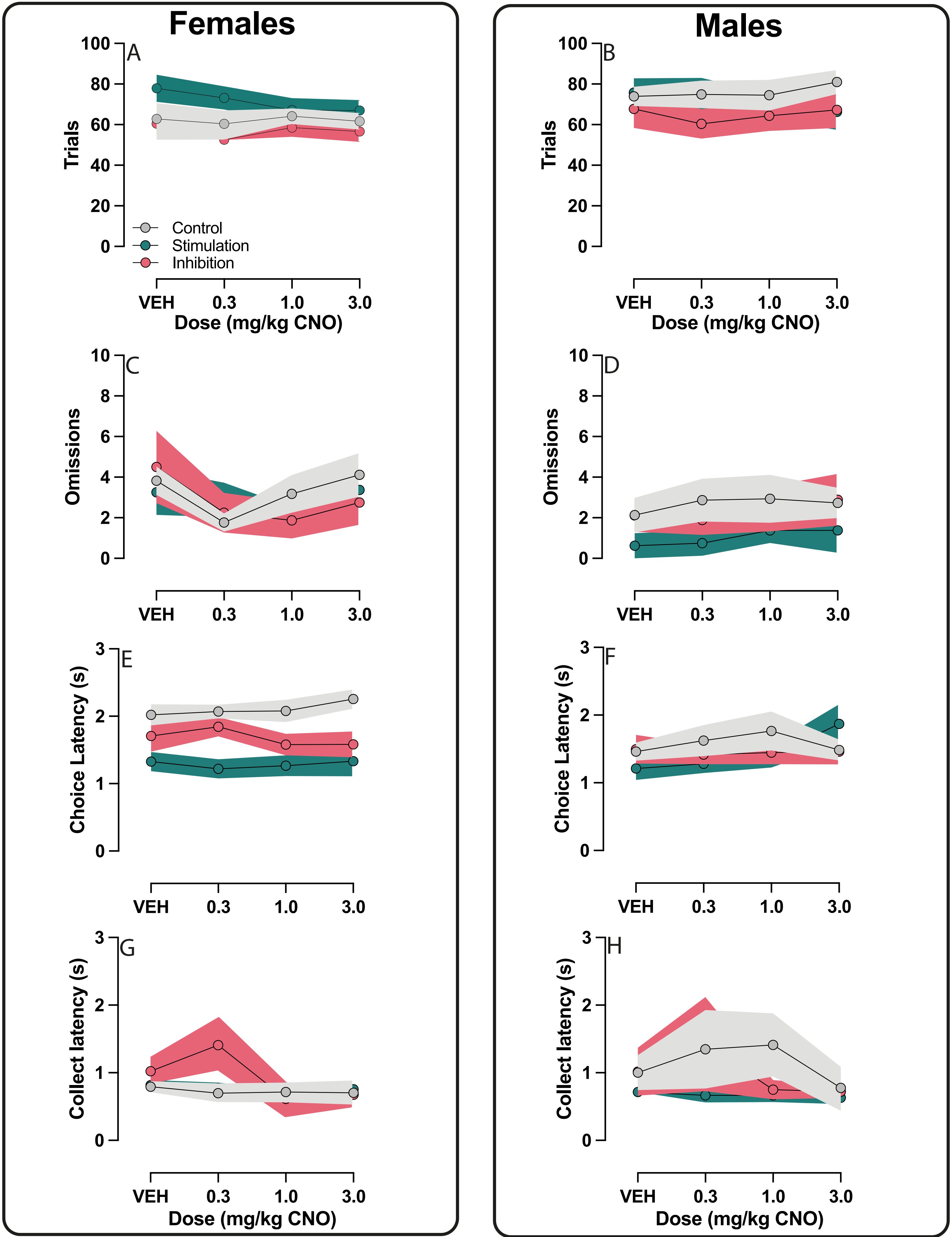

### Supplemental Figure 3

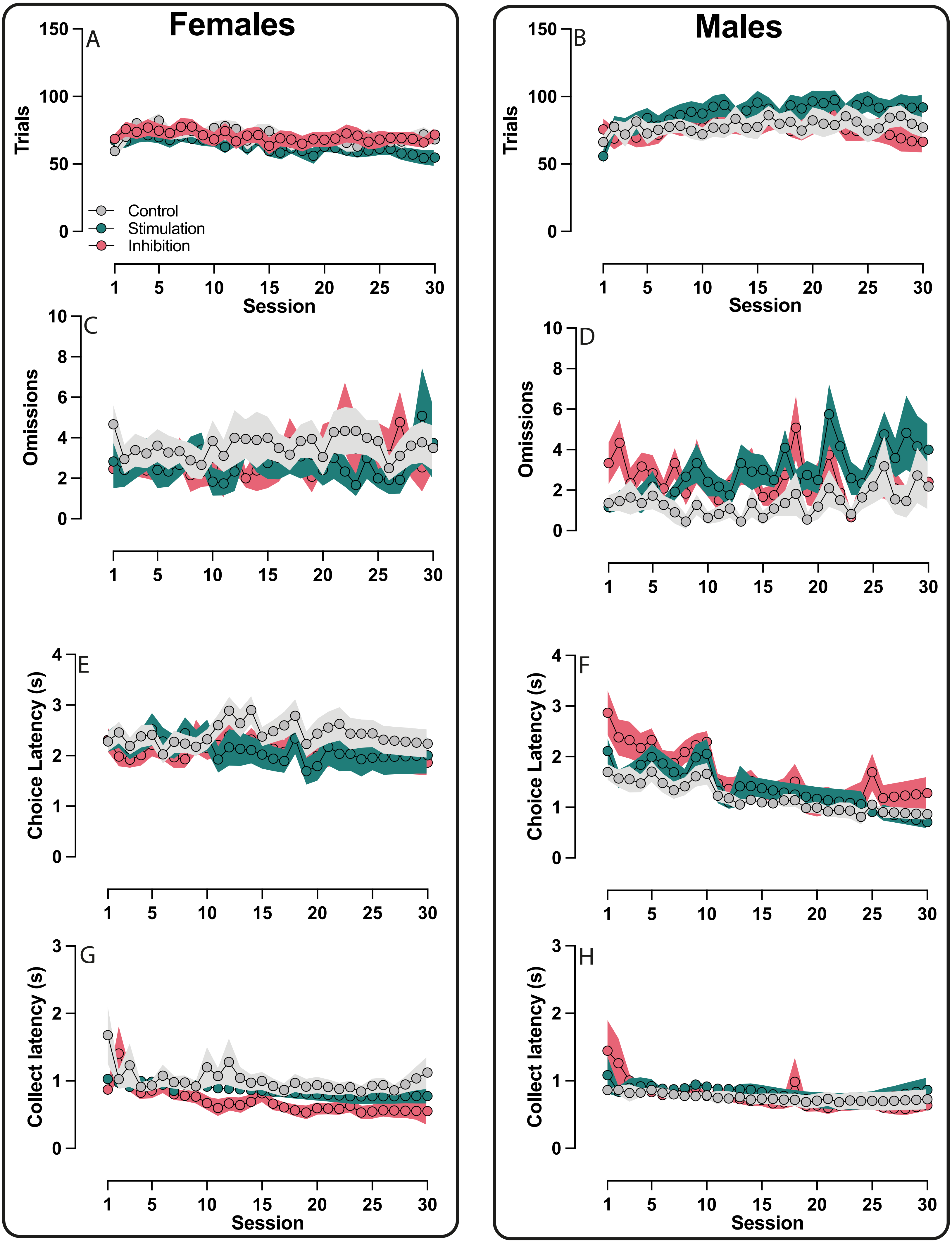
